# Epoch-specific frontal eye field control of saccades revealed by optogenetic inhibition in common marmosets

**DOI:** 10.64898/2026.11.27.769641

**Authors:** Wajd Amly, Chih-Yang Chen, Hirotaka Onoe, Kenta Kobayashi, Tadashi Isa

## Abstract

The frontal eye field (FEF) is a cortical region involved in preparation and execution of saccadic eye movements. Although neuronal activity related to saccade planning and generation has been extensively characterized in the FEF, its causal contribution to a specific epoch of saccade control remains incompletely understood. Here, we used temporally precise optogenetic inhibition in common marmosets performing saccadic tasks to examine the FEF role in saccade preparation and execution. Optogenetic inhibition at target onset, in step and gap tasks, delayed regular saccades without affecting express saccades. In contrast, inhibition during the gap period significantly reduced the frequency of express saccades and prolonged regular saccade reaction times. These effects were accompanied by changes in local field potential activity in the FEF. These findings provide causal evidence about how the FEF contributes to initiation and preparation of saccades. Our results indicate that the FEF activity during gap period contributes to preparation of express saccades, whereas FEF activity following target presentation does not. This dissociation is consistent with the hypothesis that the execution of express saccades relies predominantly on subcortical visuomotor circuits, while the FEF activity contributes to establishing the preparatory state before target onset.

## INTRODUCTION

The frontal eye field (FEF), located in the frontal cortex rostral to the arcuate sulcus in macaques, has long been recognized as a key region involved in controlling eye movements(*1*). Weak electrical stimulation of the FEF reliably elicits fixed vector contraversive saccades, suggesting its direct involvement in saccade generation(*2–5*). The FEF contains neurons with visual receptive fields, as well as neurons that discharge before or during saccadic eye and head movements(*6–10*). Anatomically, it receives convergent inputs from multiple cortical areas and projects to the superior colliculus (SC), brainstem premotor nuclei, and other oculomotor regions, placing it at the centre of the oculomotor control network(*11–13*).

Unexpectedly, however, permanent lesions(*14*, *15*), pharmacological inactivation(*16*, *17*), and cryogenic cooling(*18*, *19*) of the FEF in monkeys produce only modest and largely transient deficits. These deficits, including prolonged saccade reaction time (SRT), reduced initiation of contraversive saccades, contralesional neglect, and fixation instability, are only prominent during the acute phase and typically recover within several weeks(*20*, *21*). These transient effects may reflect functional compensation and redundancy within the distributed oculomotor network, as well as the fact that conventional manipulations alter FEF activity over relatively long timescales and therefore cannot readily distinguish its contributions at different epochs of saccade generation. In the present study, we therefore employed optogenetic inhibition of the FEF to transiently perturb neural activity with millisecond-scale temporal precision, allowing us to examine how FEF activity contributes to distinct epochs of saccade preparation and initiation (Figure 1A). This temporal precision is particularly relevant for studying express saccades, which provide a behavioural readout of the preparatory state preceding target onset. Express saccades are ultra-short-latency eye movements forming a separate distribution of SRTs from regular saccades (Figure 1B) that occur more frequently in gap task, in which a brief temporal gap separates fixation point offset from target onset, than in step saccade task, in which the target appears simultaneously with fixation point offset(*22–24*). Previous studies have shown that the occurrence of express saccades is associated with preparatory build-up activity in the SC(*25–27*). Because the FEF provides a major cortical input to the SC, it has been proposed that FEF activity contributes to this preparatory signal(*28*); however, direct causal evidence is lacking. To test this hypothesis, we examined whether optogenetic inhibition of the FEF during gap period alters the probability of express saccade generation.

**Figure 1.**
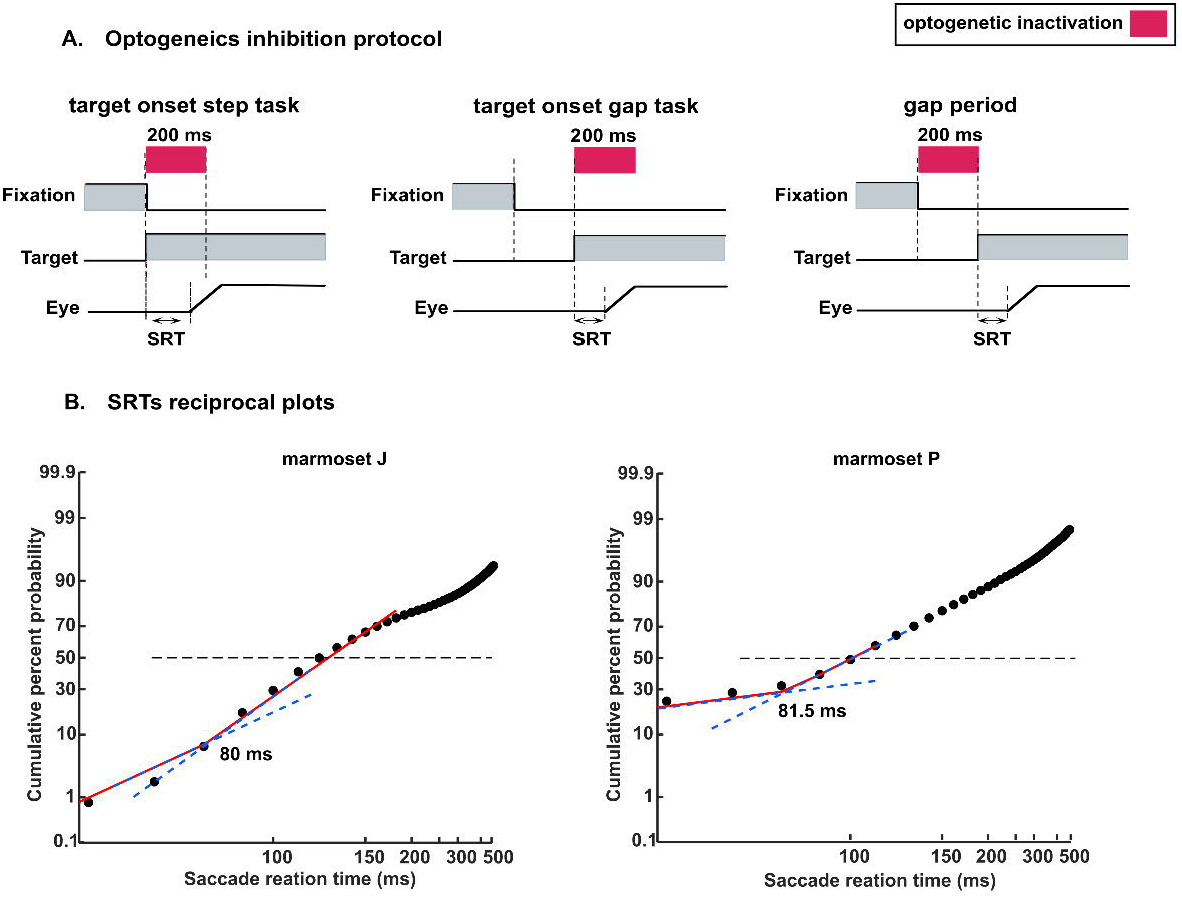
Illustration of the inhibition protocol, SRT reciprobit plot and the visual tasks. (A) Illustration of the visual tasks: in step task, the marmosets were required to make a saccade toward a visual target that appeared immediately after fixation offset. Gap task followed the same sequence, except that after a 200 ms fixation period, the fixation point disappeared for 200 ms (gap period) before the target appeared. The marmosets were required to make a saccade toward the visual target. The Pink blocks illustrate the optogenetic inactivation protocol: 200 ms illumination at target onset in step and gap tasks or 200 ms illumination during gap period. (B) Reciprobit plot showing the boundary between express and regular saccades in both marmosets.

In addition, the bimodal distribution of SRTs has led to the proposal that express and regular saccades are mediated by partially distinct visuomotor pathways(*22*). In particular, express saccades have been suggested to rely primarily on direct sensorimotor processing within the SC, whereas regular saccades depend more on cortical pathways involving the FEF. If this model is correct, inhibition of the FEF after target onset should preferentially affect regular saccades while sparing express saccades. To test this prediction, we compared the effects of temporally specific FEF inhibition using a red-shifted JAWS halorodopsin on express and regular saccades while simultaneously recording local field potentials (LFPs) from the FEF in the common marmoset (Callithrix jacchus), whose lissencephalic cortex provides unique advantages for optogenetic manipulation and electrophysiological recording.

## RESULTS

### JAWS Expression and Tet-Off–Dependent Suppression Confirmed by GFP Fluorescence

To deliver Jaws to the targeted area, we unilaterally injected an adeno-associated virus type 2.1 vector (AAV2.1-TRE3G-JAWS-KGC-GFP-ER2) together with (AAV2.1-Thy1-stTA) into the FEF of two marmosets (J and P)(*29*). In both animals, we observed robust expression in areas 8aV/6Va (Figure 2A, 2B). In marmoset P, we validated Tet-Off functionality by monitoring GFP fluorescence (Figure 2C), which was reversibly suppressed after doxycycline administration. We further confirmed the expression of JAWS-GFP in deeper layer neurons by immunostaining the post-mortem brain tissue from marmoset P, which was collected without doxycycline administration (Figure 2D).

**Figure 2.**
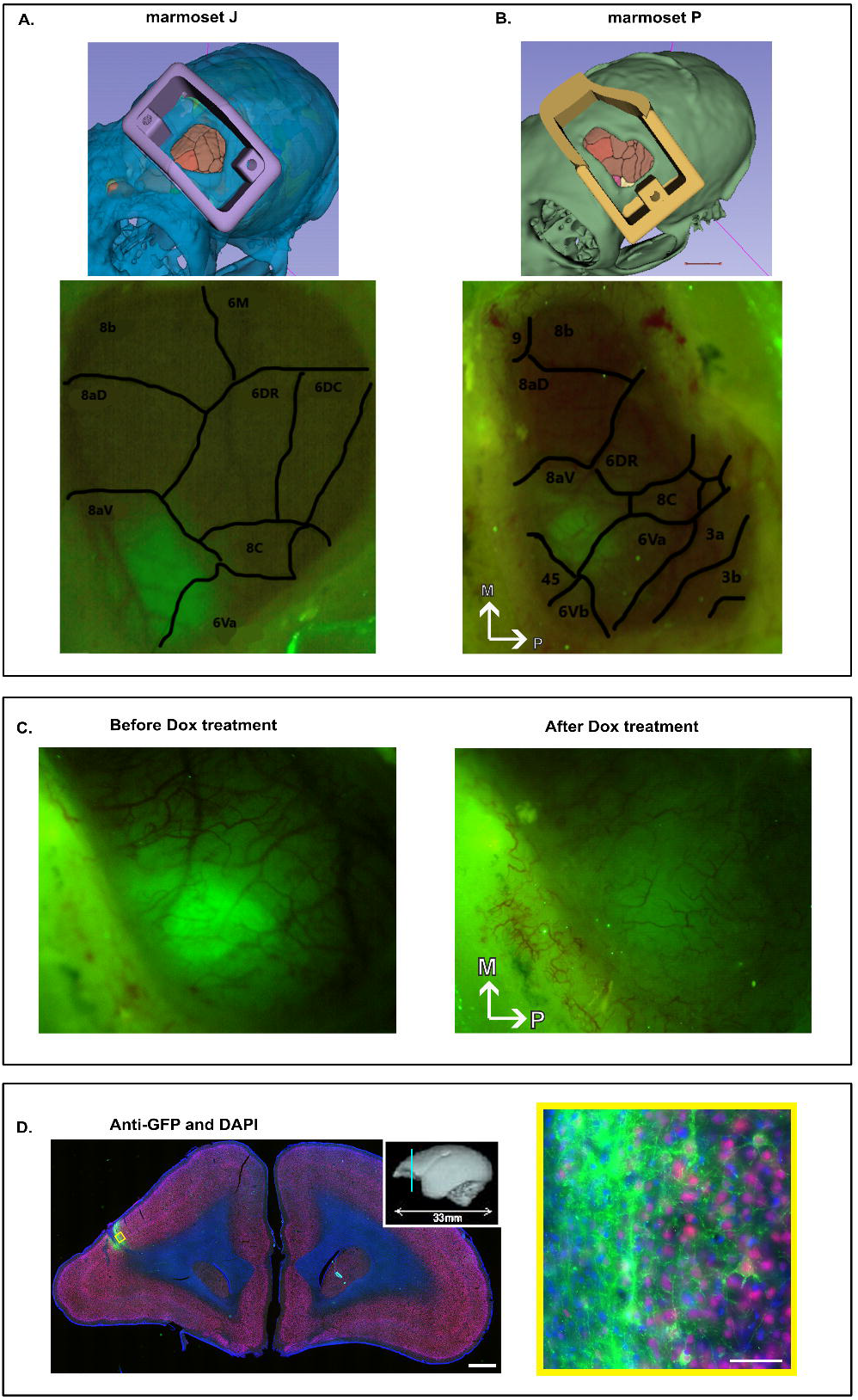
Illustration of FEF and viral expression. (A, B) Viral expression six weeks after injection, before optic fibre implantation and chamber sealing, confirmed by digital microscopy. Robust expression was observed in areas 6Va and 8aV in both marmosets. (C) Suppression of JAWS-GFP expression following doxycycline treatment was confirmed by digital microscopy. (D) Representative images showing GFP expression at the border between vector-infected and uninfected cortical area in the FEF of marmoset P. The left panel shows a low-magnification image of deeper layer cortical neurons identified by anti-NeuN immunostaining. GFP expression is shown in green, NeuN immunostaining in magenta, and DAPI staining in blue. The yellow box indicates the region enlarged in the right panel. The cyan bar in the upper-right corner of the left panel indicates the approximate anterior–posterior level of the section, located 14 mm anterior to the interaural (IA) line. The brain illustration was adapted from Stacey Jackson (CC BY-SA 4.0). Scale bars: 1 mm (left) and 50 μm (right).

### FFE Inhibition at Target Onset in Step Saccade Task

We recorded eye position in both marmosets (J and P) and LFPs in marmoset J during the step task, with optogenetic inhibition applied at target onset (Opto ON). By analysing saccades and comparing them with the corresponding OFF trials (step task trials without optogenetic inhibition), we found that the success rate of contraversive saccades was reduced in both marmosets after FEF inhibition (marmoset J: 89% to 81%, p = 0.0001; marmoset P: 75% to 68.2%, p = 0.04, Fisher’s exact test) (Figure 3A).

**Figure 3.**
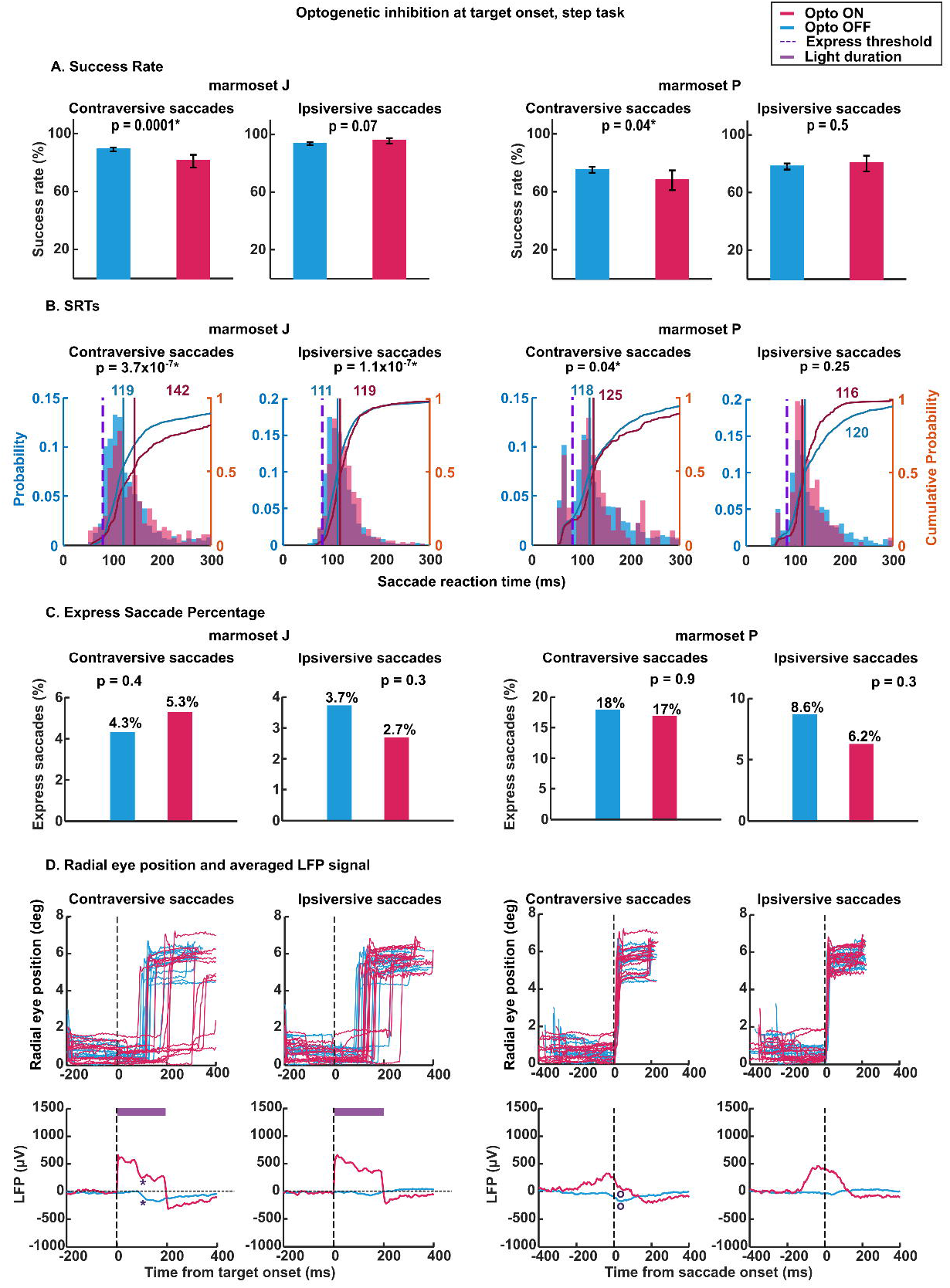
Effect of FEF inhibition at target onset in step saccade task. (A) Success rate for contraversive and ipsiversive saccades in OFF and ON trials for both marmosets. (B) SRT distributions in OFF and ON trials for contraversive and ipsiversive saccades visualized using histogram and cumulative distribution function (CDF). SRT histograms were constructed using fixed 10 ms bins and normalized to probability to allow comparison across conditions with different trial counts. Median SRT was computed for each distribution and indicated on the plots (coloured lines). The boundary between express and regular saccades is indicated by the purple line. (C) Percentage of express saccades in OFF and ON trials for each marmoset. Statistical significance is indicated with (*). (D) Radial eye position traces and corresponding averaged LFP signals aligned to target onset (left columns) for ON and OFF trials, and aligned to saccade onset (right columns). The number of trials for contraversive and ipsiversive locations under Opto ON / Opto OFF conditions is as follows: for marmoset J: 266/2129 and 451/2352, respectively; for marmoset P: 131/1336 and 177/1181, respectively.

FEF inhibition also shifted the SRT distribution rightward, resulting in prolonged SRTs median for contraversive saccades (119 to 142 ms, p = 3.7×10-7 in marmoset J, and 118 to 125 ms, p = 0.04 in marmoset P, rank-sum test) (Figure 3B). This increase in SRT was driven primarily by regular saccades (for definition of regular and express saccades, see Methods section). The median SRT of contraversive regular saccade increased significantly in marmoset J (121 to 145.5 ms, p = 8.19×10⁻9, rank-sum test) and marmoset P (132 to 135 ms, p = 0.017). In contrast, the median SRT of contraversive express saccade was not significantly affected in marmoset P (61 to 62.5 ms, p = 0.5). In marmoset J, the SRT of the contraversive express saccade showed a significant nominal decrease (72 to 66.5 ms, p = 0.03).

On the other hand, the proportion of express saccades did not differ significantly between ON and OFF trials (4.3% to 5.3%, p = 0.4 in marmoset J, and 18% to 17%, p = 0.9 in marmoset P, Fisher’s exact test) (Figure 3C). Consistent with this, a logistic regression model confirmed that the proportion of express saccades was not significantly affected by FEF inhibition at either location in marmoset J (contraversive OR = 1.24, 95% CI [0.69, 2.2], p = 0.45, FDR-adjusted p = 0.5; ipsiversive OR = 0.71, 95% CI [0.4, 1.3], p = 0.28, FDR-adjusted p = 0.4) or marmoset P (contraversive OR = 0.9, 95% CI [0.58, 1.5], p = 0.77, FDR-adjusted p = 0.8; ipsiversive OR = 0.7, 95% CI [0.37, 1.33], p = 0.3, FDR-adjusted p = 0.45). To increase statistical power and directly test whether this effect generalized across both animals, we additionally fit a mixed-effects logistic regression model pooling trial-level data from both marmosets, with a random intercept for animal to account for between-subject differences in baseline express saccade probability. The combined-animal model likewise showed no significant effect at either location (contraversive OR = 1.3, 95% CI [0.7, 2.2], p = 0.4, FDR-adjusted p = 0.5; ipsiversive OR = 0.76, 95% CI [0.4, 1.4], p = 0.35, FDR-adjusted p = 0.5).

We next examined the neural correlates of transient FEF inhibition by analysing LFP activity and radial eye position aligned to target onset (Figure 3D, left columns). Radial eye position traces in ON trials were shifted rightward relative to OFF trials, consistent with the delayed SRTs reported above. Following light onset, the LFP exhibited a prominent positive deflection, consistent with the optogenetically induced suppression of neuronal population activity. This positive deflection returned toward negative values after light offset. In addition, a negative deflection was observed approximately 120 ms after target onset in both ON and OFF trials, particularly pronounced for contraversive saccades (“*” in Figure 3D, lower left panel). To examine activity associated with movement initiation, we next aligned the LFP signal to saccade onset and plotted the trial-averaged responses (Figure 3D, right columns). A movement-related LFP deflection was observed around the time of saccade execution, with greater amplitude for contraversive saccades (“o” in Figure 3D lower right panel). To further emphasize the changes in LFP activity, we overlaid the LFP signals for contraversive and ipsiversive saccades (**Supplementary Figure 1A**). A negative deflection around saccade onset can be seen in both ON and OFF conditions for contraversive saccades.

### FFE Inhibition during Gap Period

We further collected and analysed saccades performed in the gap task when optogenetic inhibition was applied during the gap period (Opto ON) and compared them with corresponding OFF trials. Success rate of contraversive saccades was reduced in both marmosets after FEF inhibition, reaching significance in marmoset J (85% vs 80.4, p = 0.01, Fisher’s exact test) but not in marmoset P (64.6% vs 63%, p = 0.7) (Figure 4A).

**Figure 4.**
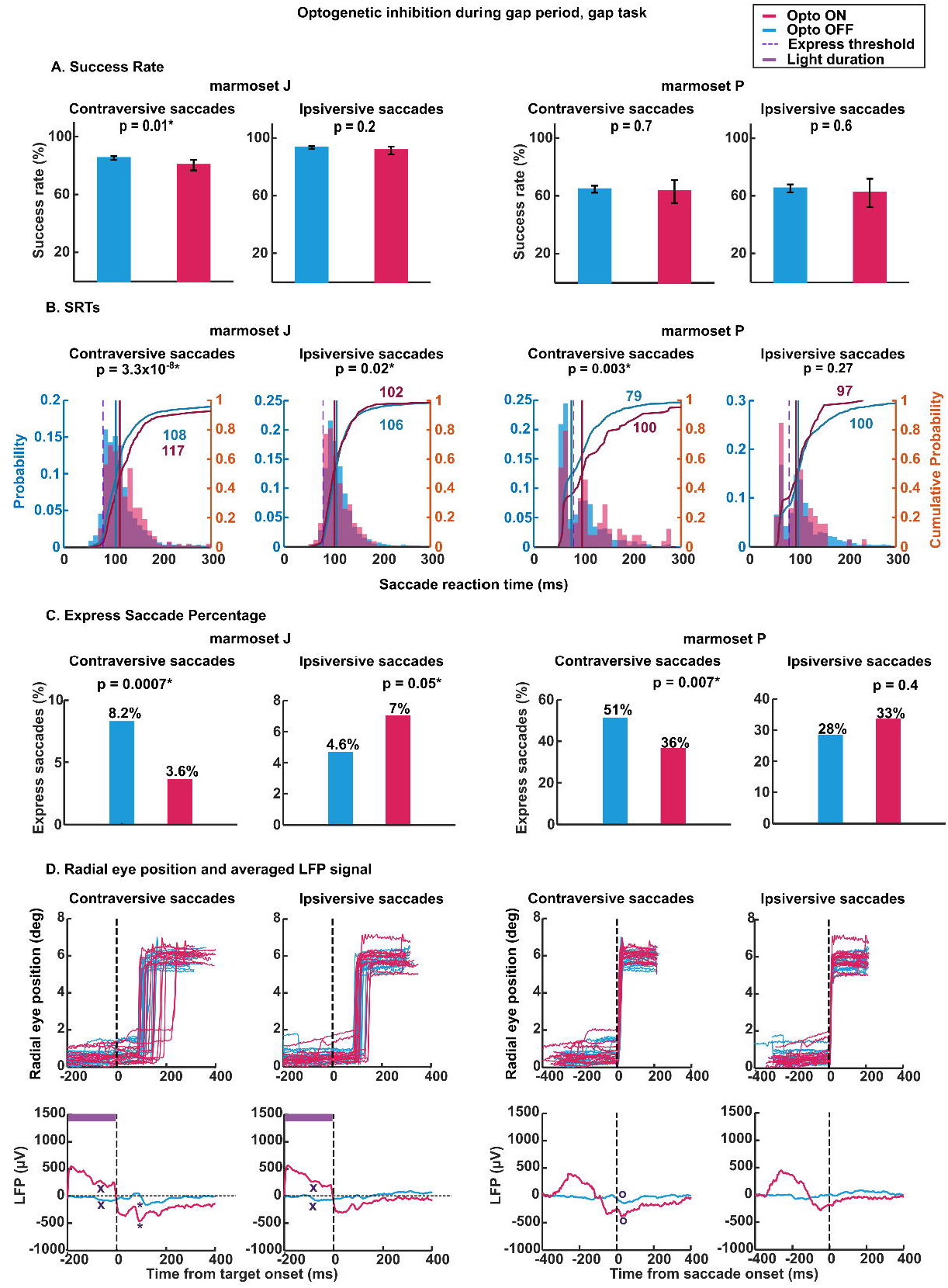
Effect of FEF inhibition during gap period. (A) Success rate for contraversive and ipsiversive saccades in OFF and ON trials for both marmosets. (B) SRT distributions in OFF and ON trials for contraversive and ipsiversive saccades visualized using histogram and CDF. SRT histograms were constructed using fixed 10 ms bins and normalized to probability to allow comparison across conditions with different trial counts. Median SRT was computed for each distribution and indicated on the plots (coloured lines). The boundary between express and regular saccades is indicated by the purple line. (C) Percentage of express saccades in OFF and ON trials for each marmoset. (D) Radial eye position traces and corresponding averaged LFP signals aligned to target onset (left columns) for ON and OFF trials. Statistical significance is indicated with (*). The number of trials for contraversive and ipsiversive locations under Opto ON / Opto OFF conditions is as follows: for marmoset J: 390/2608 and 415/2634, respectively; for marmoset P: 96/1023 and 63/782, respectively.

Furthermore, FEF inhibition during gap period significantly prolonged the median of contraversive SRTs from 108 to 117 ms, in marmoset J (p = 3.3 x 10-8) and from 79 to 100 ms in marmoset P (p = 0.003, rank-sum test) (Figure 4B). As in the previous condition, the median SRT of contraversive regular saccade increased significantly in both marmoset J (110 to 119 ms, p = 7.96×10⁻6, rank-sum test), and marmoset P (114 to 136 ms, p = 0.03), whereas the median SRT of express saccade itself was unaffected (marmoset J: 75 to 75.5 ms, p = 0.94; marmoset P: 60 to 60 ms, p = 0.3).

Although SRT of express saccade was largely unchanged, FEF inhibition during gap period markedly affected the probability of generating express saccades. The proportion of express saccades differed significantly between ON and OFF trials in both marmosets (8.2% to 3.6%, p = 0.0007 in marmoset J; and 51% to 36%, p = 0.007 in marmoset P, Fisher’s exact test) (Figure 4C). Consistent with these results, logistic regression model revealed a significant reduction in the odds of contraversive express saccades in marmoset J (OR = 0.4, 95% CI [0.24, 0.72], p = 0.0017, FDR-adjusted p = 0.02) and marmoset P (OR = 0.55, 95% CI [0.38, 0.85], p = 0.007, FDR-adjusted p = 0.04). No significant effect was observed for ipsiversive saccades in marmoset J (OR = 1.5, 95% CI [1, 2.3], p = 0.04, FDR-adjusted p = 0.1) and marmoset P (OR = 1.3, 95% CI [0.73, 2.2], p = 0.4, FDR-adjusted p = 0.5). To further assess whether the reduction in express saccade probability generalized across animals, we additionally fitted a combined mixed-effects model. This model revealed a significant reduction in the odds of contraversive express saccades (OR = 0.4, 95% CI [0.2, 0.7], p = 0.002, FDR-adjusted p = 0.012), and no effect on ipsiversive ones (OR = 1.55, 95% CI [1.02, 2.4], p = 0.04, FDR-adjusted p = 0.12).

We next examined the neural correlates of transient FEF inhibition by analysing LFP activity and radial eye position aligned to target onset (Figure 4D, left columns). Radial eye position traces in ON trials were more rightward shifted relative to OFF trials, consistent with the delayed SRTs reported above. Following light onset, the LFP exhibited a prominent positive deflection, consistent with the optogenetically induced suppression of neuronal population activity, which returned toward negative values at light offset. In addition, a negative deflection was observed approximately 50-60 ms before target onset (“**x**” in Figure 4D, lower left panels). This deflection was present in both ipsiversive and contraversive saccades and in both ON and OFF trials, presumably reflecting the preparatory activity during gap period. Following target onset, a second negative deflection was observed approximately 120 ms later, again in both ON and OFF trials, and more pronounced for contraversive saccades, presumably reflecting presaccadic activity (“*” in Figure 4D, lower left panel). To examine activity associated with movement initiation, we aligned the LFP signal to saccade onset (Figure 4D, right columns). A movement-related LFP deflection was observed around the time of saccade execution, or slightly preceding it for contraversive saccades (“**o**” in Figure 4D lower right panel), whereas no comparable deflection was observed for ipsiversive saccades. The difference in LFP activity can be clearly observed when the LFP signals for contraversive and ipsiversive saccades were overlaid (Supplementary Figure 1B). The preparatory deflection during the gap period described above was evident in both ON and OFF conditions and for both contraversive and ipsiversive saccades (the black arrows), but was absent in LFP activity during step task.

### FFE Inhibition at Target Onset in Gap Task

By analysing saccades performed in gap task under optogenetic inhibition at target onset (Opto ON) and comparing them with corresponding OFF trials, we found that FEF inhibition reduced the success rate of contraversive saccades in both marmosets, reaching statistical significance in marmoset J (88% to 80.4%, p = 0.0004, Fisher’s exact test) but not in marmoset P (79.6% vs 74.2%, p = 0.1) (Figure 5A).

**Figure 5.**
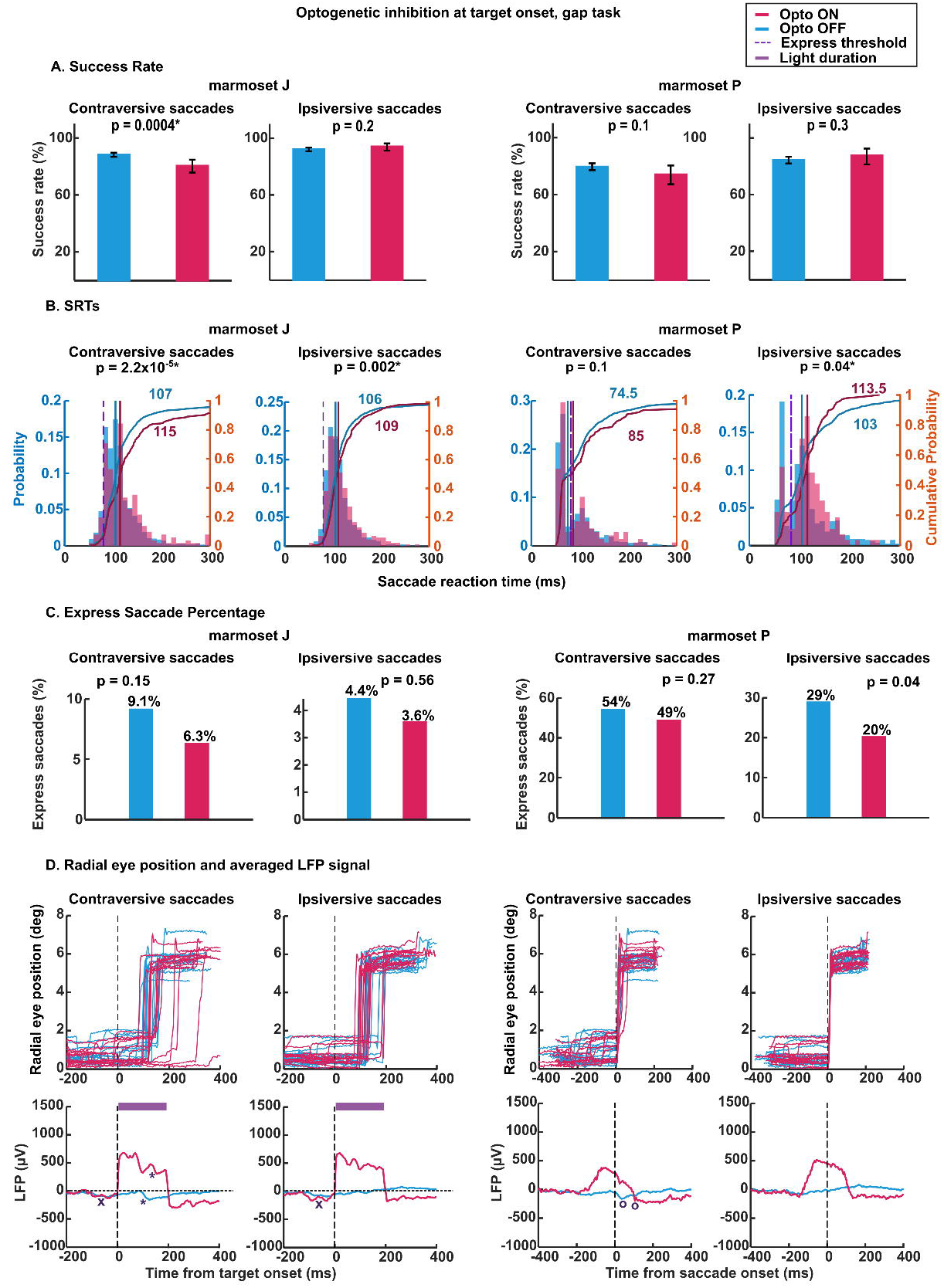
Effect of FEF inhibition at target onset in gap task. (A) Success rate for contraversive and ipsiversive saccades in OFF and ON trials for both marmosets. (B) SRT distributions in OFF and ON trials for contraversive and ipsiversive saccades visualized using histogram and CDF. SRT histograms were constructed using fixed 10 ms bins and normalized to probability to allow comparison across conditions with different trial counts. Median SRT was computed for each distribution and indicated on the plots (coloured lines). The boundary between express and regular saccades is indicated by the purple line. (C) Percentage of express saccades in OFF and ON trials for each marmoset. (D) Radial eye position traces and corresponding averaged LFP signals aligned to target onset (left columns) for ON and OFF trials. Statistical significance is indicated with (*). The number of trials for contraversive and ipsiversive locations under Opto ON / Opto OFF conditions is as follows: for marmoset J: 255/1665 and 336/1653, respectively; for marmoset P: 138/888 and 134/805, respectively.

FEF inhibition also prolonged contraversive median SRTs from 107 to 115 ms in marmoset J (p = 2.2×10-5, rank-sum test) and from 74.5 to 85 ms in marmoset P (p = 0.1) (Figure 5B). As in the other two conditions, this increase was driven by regular saccades. The median SRT of contraversive regular-saccades increased significantly in marmoset J (109 to 118 ms, p = 6.2×10-5, rank-sum test) but not in marmoset P (116 to 121 ms, p = 0.1), whereas the SRTs of contraversive express saccade was unaffected in either animal (75 to 72 ms, p = 0.4 in marmoset J; 60 to 61 ms, p = 0.8 in marmoset P).

Importantly, FEF inhibition at target onset did not significantly alter the probability of generating express saccades. Fisher’s exact tests showed no significant change in express saccade frequency (9.1% to 6.3%, p = 0.15 in marmoset J; 54% to 49%, p = 0.27 in marmoset P) (Figure 5C). Consistent with these results, a logistic regression revealed no significant location-specific optogenetic effect in marmoset J (contraversive OR = 0.66, 95% CI [0.4, 1.13], p = 0.13, FDR-adjusted p = 0.3; ipsiversive OR = 0.8, 95% CI [0.43, 1.5], p = 0.5, FDR-adjusted p = 0.5) or marmoset P (contraversive OR = 0.8, 95% CI [0.56, 1.15], p = 0.24, FDR-adjusted p = 0.45; ipsiversive OR = 0.6, 95% CI [0.4, 0.98], p = 0.04, FDR-adjusted p = 0.12). A combined-animal mixed-effects model similarly confirmed no significant effect of target-onset FEF inhibition on contraversive express saccade probability (OR = 0.63, 95% CI [0.4, 1.1], p = 0.1, FDR-adjusted p = 0.2). For ipsiversive saccades, no significant effect was observed either (OR = 0.8, 95% CI [0.4, 1.5], p = 0.5, FDR-adjusted p = 0.5). Thus, unlike FEF inhibition during gap period, inhibition at target onset prolonged regular saccade reaction time without robustly altering the probability of contraversive express saccade generation.

We next examined the neural correlates of transient FEF inhibition at target onset by analysing LFP activity and radial eye position aligned to target onset (Figure 5D, left columns). Radial eye position traces in ON trials were shifted rightward relative to OFF trials, consistent with the delayed SRTs reported above. Similar to the previous conditions, light onset elicited a prominent positive LFP deflection, consistent with optogenetically induced suppression of neuronal population activity, which returned toward negative values at light offset. In addition, a negative deflection was observed approximately 50-70 ms before target onset, during gap period (“**x**” in Figure 5D, lower left panels). This deflection was present in both ipsiversive and contraversive saccades and in both ON and OFF trials. Following target onset, a second negative deflection was observed approximately 120 ms later, again in both ON and OFF trials, and more pronounced for contraversive saccades (“*” in Figure 5D, lower left panel). To examine activity associated more directly with movement initiation, we next aligned the LFP signal to saccade onset (Figure 5D, right columns). A movement-related LFP modulation was observed around the time of execution for contraversive saccades (“**o**” in Figure 5D, lower left panel), whereas no comparable deflection was observed for ipsiversive saccades. Again, the gap period preparatory LFP signal can be clearly observed when the LFP signals for contraversive and ipsiversive saccades were overlaid (Supplementary Figure 1C). This preparatory deflection during the gap period looks similar to what was observed in the previous condition (Supplementary Figure 1B), and was evident in both ON and OFF conditions and for both contraversive and ipsiversive saccades (the black arrows).

### Greater Preparatory Activity Is Associated with Express Saccades

Motivated by our findings that gap period optogenetic inhibition selectively affected the generation of express saccades, together with the observation that FEF LFP activity during the gap period reflects a global preparatory signal, we asked whether preparatory activity differs between express and regular saccades. We first compared beta band (15–30 Hz) activity in OFF trials during gap and step tasks and found a brief peri-saccadic increase, which was more pronounced in the gap task and for contraversive location (Figures 6A, 6B). We then separated gap task saccades into express and regular saccades and found that preparatory activity was greater for express saccades (Figures 6C, 6D).

**Figure 6.**
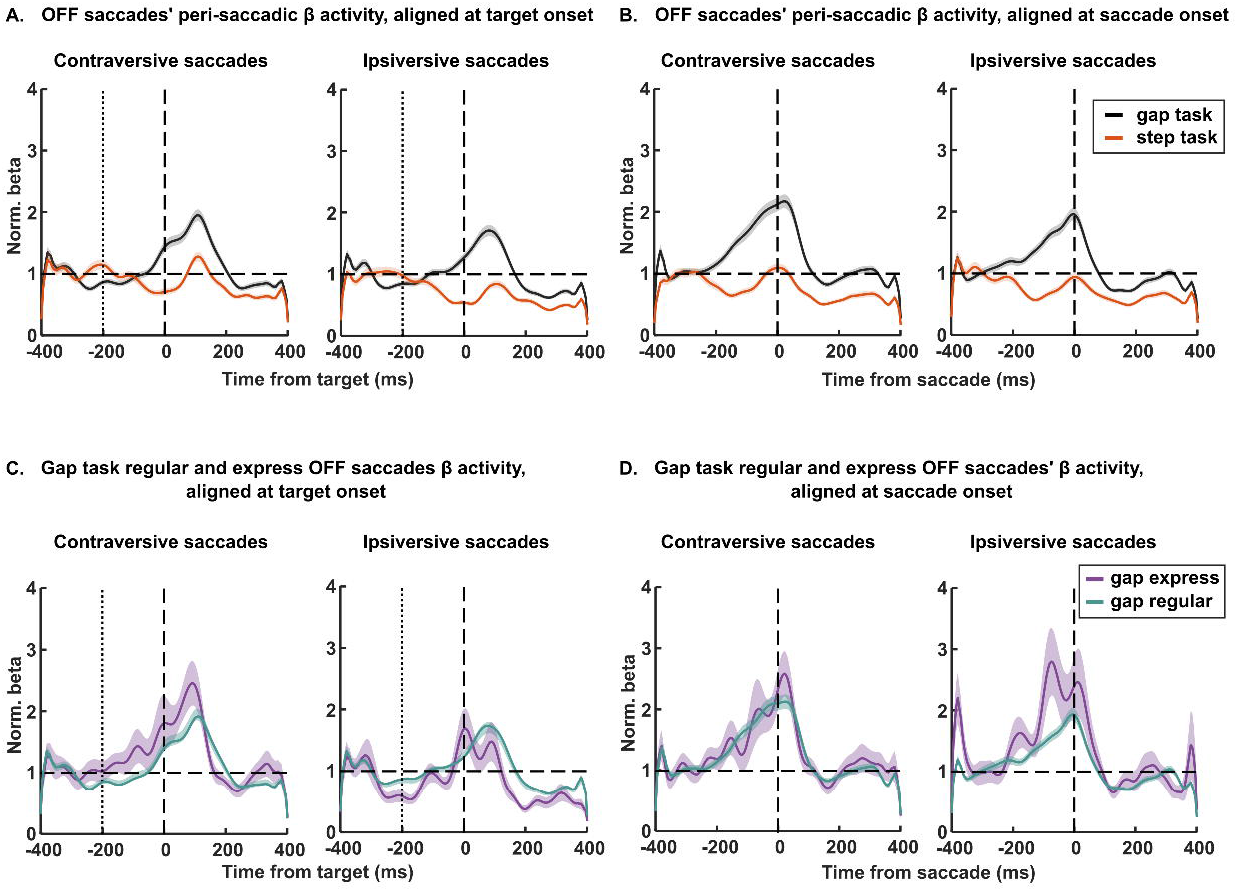
Beta band activity. (A) FEF beta activity of OFF trials aligned to target onset, and (B) aligned to saccade onset. The activity is larger in gap task than in step task. (C) FEF beta activity of regular and express saccades in gap task aligned to target onset, and (D) aligned to saccade onset. Preparatory activity is greater for express saccades.

### Suppression of JAWS Opsin Reversed Saccadic Deficits

Following doxycycline treatment and suppression of JAWS expression (Figure 2C), FEF inactivation no longer affected behavioural performance, and no significant differences were observed between ON and OFF trials. The success rate of contraversive saccades remained unchanged between OFF and ON trials (58% vs. 57.4%, respectively; p = 0.9, Fisher’s exact test) (Figure 7A). Likewise, FEF inhibition did not alter contraversive SRTs (117 vs 136 ms in OFF and ON trials, respectively; p = 0.1, rank-sum test) (Figure 7B). Logistic regression further showed no location-specific optogenetic effect on the percentage of express saccades (contraversive OR = 0.79, 95% CI [0.23, 2.8], p = 0.7; ipsiversive OR = 0.99, 95% CI [0.28, 3.5], p = 0.99), consistent with the Fisher’s exact test results (6.9% to 5.6%, p = 1) (Figure 7C).

**Figure 7.**
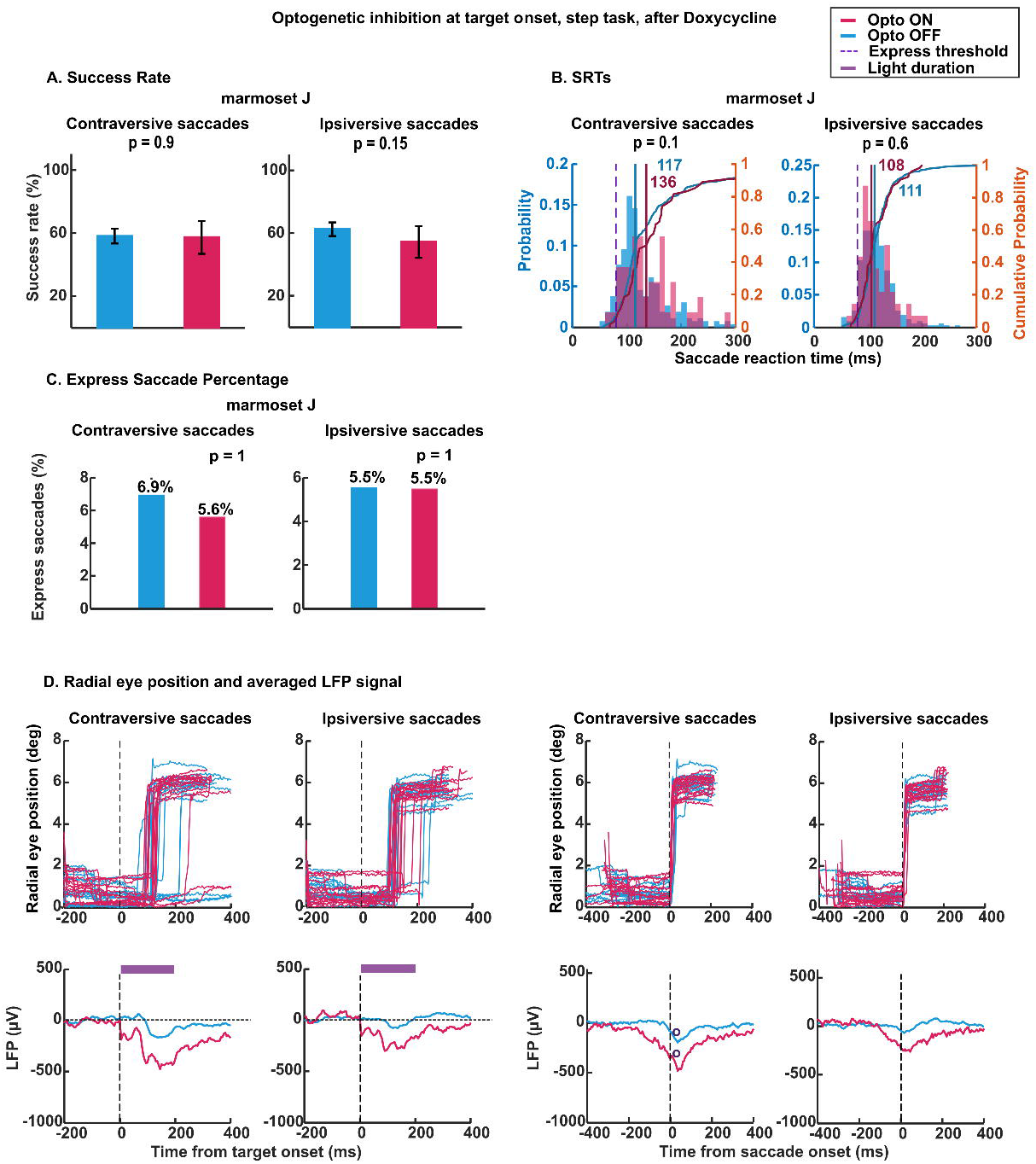
Effect of FEF inhibition at target onset in step task after doxycycline treatment. Effect in ON vs OFF trials following doxycycline treatment on (A) Success rate, (B) SRTs, (C) Percentage of express saccades, (D) Averaged LFP signals aligned at target onset and saccade onset showing comparable movement-related deflections in ON and OFF trials. FEF inactivation no longer caused significant changes in saccade behaviour or LFP signal, indicating successful silencing of JWAS-GFP expression and restoration of normal saccadic behaviour in marmoset J. SRT histograms were constructed using fixed 10 ms bins and normalized to probability to allow comparison across conditions with different trial counts. Median SRT was computed for each distribution and indicated on the plots (coloured lines). The boundary between express and regular saccades is indicated by the purple line. Statistical significance is indicated with (*). The number of trials for contraversive and ipsiversive locations under Opto ON / Opto OFF conditions is 54/261 and 55/309, respectively.

The absence of behavioural effects was accompanied by comparable LFP dynamics between ON and OFF trials. The LFP signal recorded during ON trials exhibited a negative deflection similar to that observed during OFF trials, although with a greater amplitude (Figure 7D). Moreover, the movement-related LFP deflection occurred at similar time points in both conditions for contraversive saccades (“o” in Figure 7D, lower left panel), while radial eye position traces were nearly identical between ON and OFF trials. Together, these behavioural and electrophysiological findings confirm that doxycycline-mediated suppression of JAWS expression effectively abolished the effects of optogenetic FEF inhibition.

## DISCUSSION

In the present study, we investigated how transient inhibition of the right FEF influenced saccade metrics and LFP dynamics when applied at three distinct epochs of saccade preparation: at target onset in step task, during the gap period, and at target onset in gap task. The timing of FEF inhibition produced distinct behavioural effects, suggesting that the contribution of the FEF to saccade generation likely depends on the stage of visuomotor processing at which its activity is disrupted.

At the neural level, transient optogenetic inhibition of the FEF revealed dissociable preparatory and movement-related LFP dynamics (Figures 3D, 4D, 5D; Supplementary Figures 1A, 1B, 1C). During the gap period, a spatially non-selective negative deflection emerged approximately 50-70 ms before target presentation. Given that the upcoming target direction was not yet known to the animal at this time, and that target was presented with equal probabilities and strength in the left and right visual fields, the presence of this deflection for both ipsiversive and contraversive saccades suggests that it is not strongly directionally tuned, and may instead reflect a global change in network state associated with fixation disengagement and increased oculomotor readiness, consistent with reduced fixation-related inhibition during gap effect(*30*, *31*), and non-spatially selective FEF activity following fixation offset(*32*),(*33*). Following target onset, a second negative deflection emerged approximately 120 ms later and was more pronounced for contraversive saccades. This component was preserved across the three inhibition conditions and coincided temporally around saccade onset, suggesting a movement-related population dynamics linked to saccade initiation rather than purely sensory encoding, consistent with previous LFPs in the FEF(*34*),(*35*, *36*). To further characterize this preparatory activity, we examined beta-band activity across task conditions and saccade types. Beta-band activity was stronger during gap task (Figures 6A, 6B), consistent with enhanced preparatory activity during the gap period. Moreover, within the gap task, beta-band activity was greater for express saccades than for regular saccades (Figures 6C, 6D), suggesting that stronger preparatory activity is associated with the generation of express saccades, consistent with studies in macaque monkeys(*27*). Thus, the LFP findings provide causal evidence that the FEF regulates saccadic behaviour primarily by modulating the preparatory network state of the oculomotor network and the timing of movement initiation, and link the level of preparatory FEF activity to the occurrence of express saccades, raising the question of whether reducing this preparatory activity would also alter express saccade generation at the behavioural level.

At the behavioural level, the proportion of contraversive express saccades was reduced when the FEF was inhibited during gap period (Figure 3C) but not when inhibition was applied at target onset (Figures 4C, 5C). This reduction was statistically significant in both marmosets. Consistent with previous studies of reversible FEF inactivation in macaques, express saccades were reduced but not abolished(*37*). The persistence of some express saccades suggests that sufficient preparatory activity in the SC can still develop in the absence of normal FEF input(*18*). The gap paradigm places the oculomotor system in a highly preparatory state because fixation offset precedes target presentation. This state may increase the dependence of rapid saccade generation on FEF activity. Our findings therefore suggest that FEF activity contributes to maintaining the preparatory state that facilitates express saccade generation under conditions of elevated motor readiness, rather than directly determining the timing of express saccades once the target appears. Consistent with this interpretation, FEF inhibition applied at target onset did not significantly alter either the latency or the proportion of express saccades. Thus, the effects of FEF inhibition appear to depend on the temporal epoch at which the perturbation occurs, with disruption during the preparatory gap period having a greater impact on express saccade generation than disruption at target onset. The relative preservation of express saccade latency following FEF inhibition at target onset is also consistent with the possibility that the final execution of these very rapid responses relies heavily on downstream short loop brainstem circuits. One proposed mechanism is a short latency pathway through the SC, potentially involving transmission from its superficial visual layers to deeper visuomotor layers(*38*, *39*). However, the present results do not directly establish the contribution of this pathway. Simultaneous recordings or pathway-specific manipulation of SC neurons during FEF inhibition would therefore be essential to determine whether the preserved express saccade latency reflects the recruitment of such a subcortical pathway.

Because express saccades represent the fastest end of the SRT distribution, the reduction in their proportion suggested that FEF inhibition may affect saccade initiation timing more generally. We therefore examined the effects of FEF inhibition on SRT. FEF inhibition commonly prolonged contraversive SRTs across the three experimental conditions, although the magnitude and statistical significance of the effect differed between animals and conditions (Figures 3B, 4B, 5B). Importantly, this effect was predominantly associated with regular saccades. The prolongation of SRTs is consistent with previous studies using reversible FEF inactivation in macaques, which reported longer SRTs for contraversive saccades with substantial overlap between inactivation and control distributions(*16*, *40*, *41*). The preservation of express saccade latency despite the prolongation of regular saccade latency further suggests that the FEF may modulate the probability or readiness of saccade initiation without directly determining the intrinsic timing of the fastest saccades.

Interestingly, ipsiversive saccades in marmoset P exhibited shorter SRTs following FEF inhibition. A similar, although weaker, facilitation was observed in marmoset J when inhibition was applied during the gap period. The direction-dependent changes in SRT may reflect a redistribution of activity within the saccade-related population, whereby suppression of neurons tuned to one direction could alter the relative balance of activity across direction-selective populations. Such an effect would be consistent with population coding frameworks, in which motor commands emerge from the weighted activity of directionally tuned neuronal populations(*42–46*). Alternatively, because the FEF is embedded within a bilateral oculomotor network, inhibition of one FEF may alter the balance of activity between the two hemispheres and thereby influence saccades in the ipsiversive direction. Additionally, FEF inactivation has been reported to reduce executive control or gating mechanisms, occasionally leading to premature or less constrained saccades(*47*, *48*). However, these interpretations remain speculative, and the ipsiversive SRT changes should be interpreted cautiously, while the more consistent prolongation of contraversive SRTs provides stronger evidence for a role of the FEF in saccade initiation.

In addition, our results showed that transient inactivation of the right FEF significantly reduced the success rate of contraversive saccades, although the magnitude and statistical significance of this effect varied across animals and experimental conditions (Figures 3A, 4A, 5A). Overall, this pattern is consistent with the established role of the FEF in facilitating the initiation and execution of contraversive saccades(*1*, *12*, *49*), and may reflect a reduction in the effective excitatory drive available to the saccade-generating network(*37*, *40*, *50*).

Finally, following doxycycline treatment, JAWS expression was suppressed, as confirmed by the reduction in GFP fluorescence (Figure 2C), and the behavioural effects were no longer observed (Figure 7). The associated LFP dynamics were also largely similar between conditions; however, its negative deflection remained slightly larger in amplitude. This residual negativity could reflect a light-evoked retinal signal that persists independently of opsin expression, consistent with reports that optical fibre illumination can activate endogenous retinal photoreceptors and generate downstream cortical responses even without viral transduction(*51*). Alternatively, it could reflect a light-induced artefact related to the photoelectric effect(*47*). Together, this dissociation between opsin-dependent and light-dependent LFP components, combined with the full behavioural recovery following doxycycline, supports the interpretation that the observed behavioural impairments were primarily mediated by JAWS-dependent inhibition of the FEF, reflecting targeted and controllable modulation of cortical excitability, while the remaining LFP component may include light-dependent effects unrelated to opsin-mediated inhibition.

## MATERIALS and METHODS

### Animal Preparation and Surgical Procedure

Two male adult marmosets born in the breeding colony at the Kyoto University Animal Research Facility, aged 3-4 years, participated in this study: marmoset J and marmoset P. The experiments were conducted following the guidelines of the Japan Neuroscience Society and the Science Council of Japan and were approved by the Animal Ethics Committee at Kyoto University, Japan.

For this experiment, the marmosets underwent three separate surgeries under aseptic conditions. In the first surgery, a custom-designed headpost was mounted on each marmoset, tailored to the individual’s skull based on CT scan reconstruction. The headpost was 3D-printed using Rigid 4000 Resin (Form2, Formlabs, U.S.A.) and attached to the skull with Super-Bond (Sun Medical Co., Ltd., Japan). In the second surgery, a craniotomy covering FEF was performed and the viral vector was injected. The craniotomy location targeting the FEF was planned by aligning each marmoset’s MRI with a reference marmoset brain from the Brain/MINDS data portal(*52*). The FEF injection locations were selected based on microstimulation mapping within the cranial window, which evoked contraversive saccades at low current intensity(*53*, *54*). In the third surgery, and after confirming the virus expression, a TeleOpt-ELG fibre (diameter 500 μm, length 3 mm, Bio Research Centre, Co., Ltd., Japan) accompanied by two LFP recording electrodes (length 1.1 mm from the fibre tip) was implanted above the infected area in marmoset J. A TeleOpt fibre (diameter 500 μm, length 3 mm, Bio Research Centre, Co., Ltd., Japan) was implanted above the infected area in marmoset P. Both fibres were fixed with Super-Bond (Sun Medical Co., Ltd., Japan) and protected by attaching a 3D-printed chamber that covers them. All surgical procedures and postoperative care were performed as previously described(*55*, *56*).

### Step and Gap Saccade Tasks and Monitor Setup

The monitor setup used for stimulus presentation was the same as described previously(*56*). The step saccade task required the animal to make a saccade toward a target that appeared immediately after the fixation offset (Figure 1A). Each trial began with a 200 ms fixation period, after which the target location was randomly selected from one of the four oblique positions. The gap saccade task had the same configuration, except that a 200 ms gap period was inserted between the fixation offset and target onset (Figure 1A). Target eccentricity was fixed at 6° for marmoset J, and 4° for marmoset P. Successful responses were rewarded with 0.05 ml of a solution prepared by mixing baby supplement banana pudding (Kewpie Corp., Japan) with banana-flavoured Mei-balance (Meiji Holdings Co., Japan). One session of the task consisted of 24 trials. On average, the marmosets completed seven sessions per day, totalling 168 trials. Across the entire experiment, we collected 39,910 trials from marmoset J and 21,399 trials from marmoset P.

### Eye Tracking, Calibration and Saccade Detection

Eye movements were recorded using the EyeLink 1000 Plus system (SR Research, Canada) at a sampling frequency of 500 Hz. Both pupil and corneal reflection signals were used to ensure better accuracy. Eye position calibration and validation were performed following the procedure described in Chen et al. 2021(*57*). Saccades were detected using U’n’Eye(*58*), which has been previously applied to marmoset saccade detection(*55*, *56*). Subsequent analyses included only contraversive upper-left and ipsiversive upper-right saccades.

### Express Saccade Identification and SRT Calculation

SRT was defined as the interval between target onset and saccade initiation. Following the described method in Amly et al. (2024)(*56*), anticipatory saccades (SRT < 50 ms) were excluded from the analysis. To distinguish express from regular saccades, SRTs were analysed using the reciprobit method(*59*). Briefly, SRTs were transformed to their reciprocals (1/SRT) and plotted against cumulative probability on a probit scale. A two-segment piecewise linear regression was then fitted to the resulting distribution using a least-squares curve fitting approach. The hinge point of the fitted model, converted back to milliseconds, was taken as the boundary between express and regular saccades. This analysis yielded thresholds of 80 ms for marmoset J and 81.5 ms for marmoset P (Figure 1B). After all selection and exclusion criteria were applied, 16,546 and 7,224 trials from marmoset J and P, respectively, were included in the analysis.

### Logistic Regression

To assess whether optogenetic inhibition altered the probability of generating an express saccade, trials were classified as express or regular based on a fixed SRT threshold as described above (Marmoset J: 80 ms; marmoset P: 81.5 ms, Figure 1B). For each animal, a generalized linear model with a binomial distribution and logit link was fit to trial-level data:

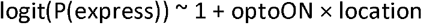

where optoON was a binary indicator of optogenetic condition (1 = ON, 0 = OFF) and location was a categorical predictor coding target direction (ipseversive vs. contraversive relative to the inactivated hemisphere). Including the optoON × location interaction allowed the effect of optogenetics to vary by location rather than assuming a single fixed effect across the visual field. Model coefficients were exponentiated to yield odds ratios (OR), where OR > 1 indicates increased odds of an express saccade under opto ON relative to OFF, and OR < 1 indicates decreased odds. Because the model included an interaction term, the OR for opto at each individual location was computed as a linear combination of the relevant coefficients (main effect of optoON, plus the corresponding interaction term where applicable), with standard errors and 95% confidence intervals derived from the model’s full coefficient covariance matrix (Wald approximation). Overall model fit was assessed via a likelihood-ratio chi-square test comparing the fitted model to an intercept-only (constant) model. Because location-specific opto effects were assessed across three inhibition timing conditions, two locations, and two animals (12 comparisons total), p-values were corrected for multiple comparisons using the Benjamini-Hochberg false discovery rate procedure (FDR) (Q = 0.05) to get FDR-adjusted p.

To evaluate whether location-specific opto effects on express-saccade probability generalized across animals, we additionally fit, separately for each inhibition condition, a mixed-effects logistic regression model pooling trial-level data from both marmosets:

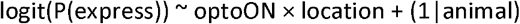

with a random intercept for animal to account for between-subject baseline differences in express-saccade probability. Because the two animals were tested at different target eccentricities (4° and 6°), animal identity and eccentricity were fully confounded in this combined dataset, and the random intercept should be interpreted as capturing between-subject variability broadly rather than an eccentricity-independent animal effect. These combined-animal models were treated as supplementary robustness checks rather than additional pre-specified location-specific comparisons, and were not included in the 12-test multiple-comparisons correction applied to the per-animal analyses.

### Optogenetic Inactivation Protocol

For optogenetic inactivation, red light (630 nm) was delivered above the FEF with peak irradiances of approximately 60 mW/mm² and 28 mW/mm² in marmoset J and P, respectively, measured at the tip of the optical fibre(*47*, *60*). Inhibition was applied for 200 ms on every fifth trial, either at target onset or during gap period (Figure 1A), to prevent heating of the cortex(*47*). Trials with optical inhibition were referred to as ON trials, whereas trials without optical inhibition were referred to as OFF trials.

### Viral Vector Injection

For the optogenetic experiment, excitatory neurons in the right FEF were selectively targeted by two unilateral injections of the viral vector delivered perpendicular to the coronal plane into area 8aV at a depth of 1.4 mm below the cortical surface. Each animal received a total volume of 1.6 µL (0.8 µL per injection) of a red-shifted JAWS halorhodopsin construct (AAV2.1-TRE3G-JAWS-KGC-GFP-ER2) combined with AAV2.1-Thy1-stTA (mixture titer: 2 × 10⁹ vg/µL). The viral vector was infused at a rate of 0.1 µL/min for 8 minutes. Following each injection, the needle was left in place for 5 minutes before slow retraction to minimize backflow of the viral solution along the injection tract. Around six weeks were allowed for the expression of JAWS before proceeding with the next stage of the experiment. Viral expression was confirmed by visualizing GFP fluorescence in the targeted region using a digital microscope (Dino-Lite, Co., Ltd., USA).

### LFP Recording and Alignment with Optogenetic Stimulation

LFP signals were recorded at 200 Hz (Nyquist frequency 100 Hz), the maximum sampling rate supported by the recording device, and segmented relative to the timing of optogenetic stimulation. The DIO signal corresponding to light stimulation was used to identify the onset and offset of each light pulse, allowing alignment of LFP activity to the optogenetic inactivation period.

### LFP Beta Band Analysis

Instantaneous beta power was obtained from a zero-phase Butterworth band-pass filter followed by the squared Hilbert envelope. For each condition and target location, trial-averaged power was divided by the mean of that average in the −350 to −250 ms window of the plotted alignment, so that a value of 1 is the mean power in that baseline. Delta (1–4 Hz) has a period comparable to the ±400 ms analysis window, so Hilbert power in that band is not reliable, and high-gamma (50–90 Hz) lies against the 100 Hz Nyquist limit of our 200 Hz sampling and cannot resolve a spike-related high-gamma burst.

### Doxycycline Treatment

To suppress JAWS expression, doxycycline (2 mg/mL) was administered orally twice daily, incorporated into Castella cake as a palatable carrier. In marmoset J (TeleOpt-ELG), doxycycline treatment was used to evaluate behavioural performance and LFP activity during JAWS suppression. In marmoset P (TeleOpt), doxycycline treatment was used to validate the Tet-regulated functionality by confirming doxycycline-dependent suppression and subsequent recovery of JAWS expression after doxycycline withdrawal prior to sealing the FEF chamber.

### Histological Assessment and Anti-GFP Immunohistochemistry

After completion of the behavioural experiment, marmoset P was deeply anesthetised with ketamine followed by sodium thiopental (25[mg/kg, i.p.) and transcardially perfused with 0.1 M phosphate-buffered saline (PBS), followed by 4% paraformaldehyde (PFA) in PBS. We then extracted the whole brain and post-fixed it overnight in 4% PFA. After cryoprotection in 30% sucrose in PBS, the brain was sectioned coronally at 40-μm thickness using a freezing microtome.

Brain sections were processed for immunohistochemistry using antibodies against GFP and NeuN, with DAPI counterstaining. Briefly, sections were incubated with a rabbit anti-GFP antibody (1: 5,000; A11122, Invitrogen), followed by an AF4880-conjugated goat anti-rabbit IgG secondary antibody (1: 200; A-11034). For NeuN immunostaining, sections were incubated with a mouse monoclonal anti-NeuN primary antibody (MAB377, Millipore), followed by an AF647-conjugated goat anti-mouse IgG (1: 200; ab150115, Abcam). Sections were mounted using VECTASHIELD Antifade Mounting Medium with DAPI (H-1200; Vector Laboratories, Newark, CA, USA). Images were acquired using a Keyence X-810 fluorescence microscope (Keyence, Japan).

## ACKNOWLEDGMENTS

This work was supported by the Japan Society for the Promotion of Science (JSPS) Grants-in-Aid for Scientific Research (Grant Nos. 23H03700 to C.C. and 25K24531 to T.I.); the Brain/MINDS program of the Japan Agency for Medical Research and Development (AMED; Project No. 19dm0207093h0001 to H.O.); Brain/MINDS 2.0 (AMED; Project No. 24wm0625201h0001 to T.I.); AMED Grant No. JP23wm0625001 to K.K.; and AMED Moonshot Goal 7 (Project No. 24zf0127010h0001 to T.I.). We thank Kaoru Isa and Aki Tanimizu for performing the histological analyses.

## AUTHOR CONTRIBUTIONS

The experiments were designed by W.A., C.C. and T.I. W.A. conducted the experiments, surgeries, behavioural training, data collection and analysis, and drafted the manuscript. C.C. helped with the surgeries and provided manuscript editing and revisions. H.O. conducted the MRI scans. K.K. designed and produced the viral tools. T.I. supervised the project and approved the final manuscript. All authors contributed to and approved the submitted version.

## FIGURE LEGENDS

**Supplementary Figure 1. Averaged, overlaid LFP activity.**

(**A**) Averaged overlaid LFP signals for contraversive and ipsiversive saccades, aligned to target onset (left columns) and saccade onset (right columns), during step task with optogenetic inhibition applied at target onset. **(B)** Averaged overlaid LFP signals for contraversive and ipsiversive saccades, aligned to target onset (left columns) and saccade onset (right columns), with optogenetic inhibition applied during the gap period. (C) Averaged overlaid LFP signals for contraversive and ipsiversive saccades, aligned to target onset (left columns) and saccade onset (right columns), during gap task with optogenetic inhibition applied at target onset.

