## Supplementary figures and images for "Epoch-specific frontal eye field control of saccades revealed by optogenetic inhibition in common marmosets"

### Supplementary figure 1

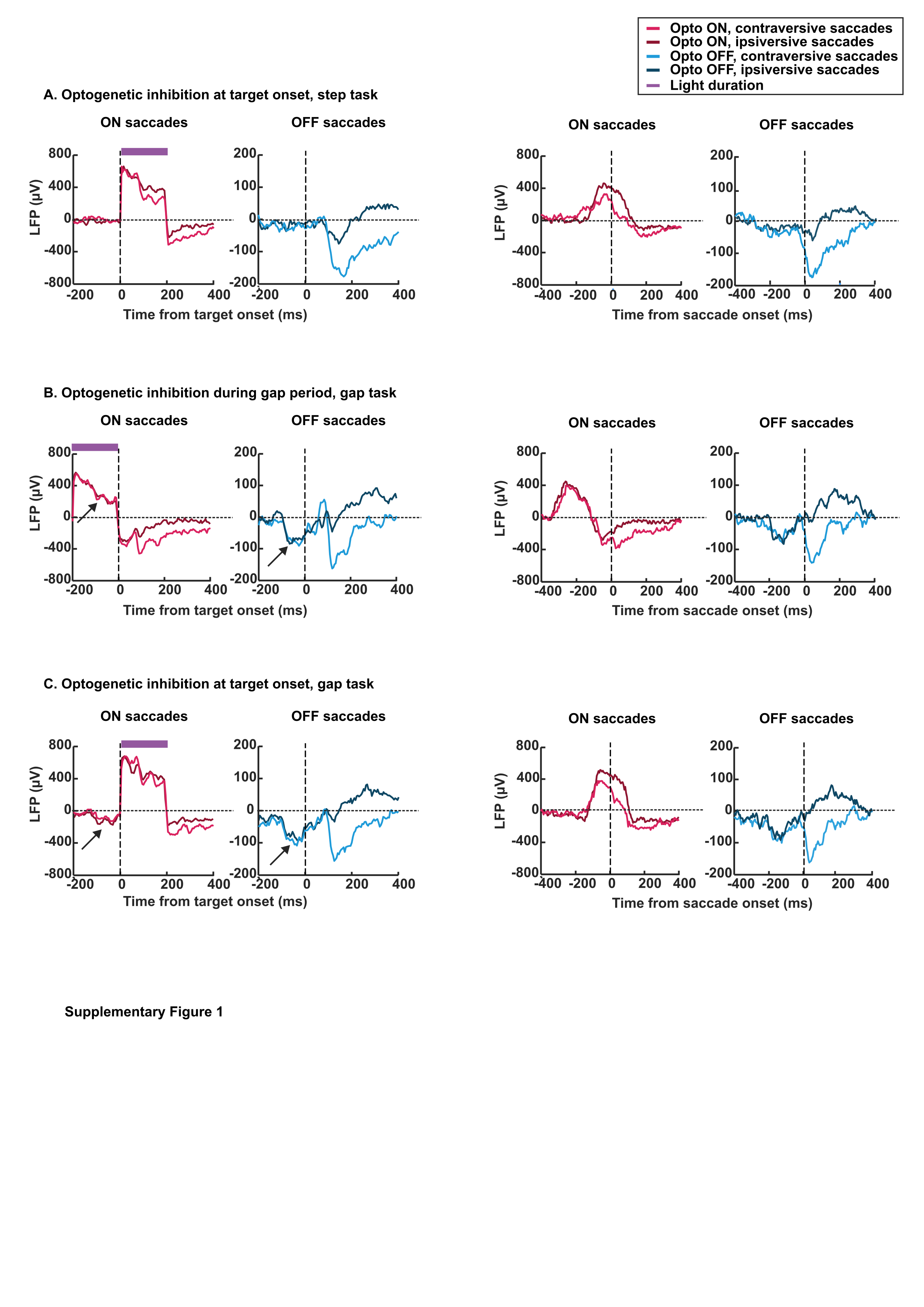
